# Tongue bite apparatus highlights functional innovation in a 310-million-year-old ray-finned fish

**DOI:** 10.1101/2025.05.10.653277

**Authors:** Sam Giles, Matthew A. Kolmann, Matt Friedman

## Abstract

Gill skeleton modifications for processing prey represent a major source of functional innovation in living ray-finned fishes. Here we present the oldest actinopterygian tongue bite, derived from the gill skeleton, in the Early Pennsylvanian (∼310 Ma) †*Platysomus parvulus*. Unrelated to extant tongue biters, this deep-bodied taxon possesses a large, multipartite basibranchial tooth plate opposing an upper tooth field centered on the vomer. This branchial structure occurs in conjunction with toothed jaws, indicating a role for both in feeding. †*P. parvulus* illustrates the assembly of the tongue bite in the geologically younger †Bobasatraniidae: large opposing dorsal (vomerine) and ventral (basibranchial) crushing plates associated with toothless jaws. The origin of tongue bites falls within the Carboniferous actinopterygian radiation, although postdates the first signs of durophagy in other ray-finned lineages by several million years. This lends support to a protracted ‘long-fuse’ model of actinopterygian diversification in the aftermath of the End-Devonian Mass Extinction.

## 1. Background

Living ray-finned fishes display a remarkable range of specializations associated with feeding. Among the most celebrated of these adaptations are modifications to the branchial apparatus in teleosts [1, 2], including so-called “tongue bites”, which involve opposing dentition on the ventral surface of the braincase and palate and tooth plates on the dorsal surface of the median gill skeleton [3, 4]. Tongue bites and modified pharyngeal jaws serve as additional zones for prey processing, with their decoupling from mandibular jaws hypothesized to increase functional versatility [5]. Extant teleost lineages with tongue bites are relatively young, with records extending only to the Mesozoic [6]. This is a time during which feeding innovations among fishes and other predatory taxa have been implicated in major macroecological shifts in aquatic communities [7].

While the Mesozoic represents a key evolutionary interval for many extant groups, entirely extinct fish lineages show examples that occurred independently of—and geologically long before—those in living species. Carboniferous fossils record the first episode of major morphological divergence among actinopterygians [8], apparent in external features like body shape and dental structure [9, 10]. Information about gill arches, in contrast, requires uncrushed specimens examined via acid preparation [11] or computed tomography [12]. The few described Carboniferous gill skeletons belong to taxa with conservative cranial and postcranial anatomy [12, 13], and do not differ substantially from those of Devonian examples [14].

Here we report gill-arch structure in a uniquely three-dimensional skull of the late Carboniferous actinopterygian †*Platysomus parvulus* (“†” precedes extinct taxa; [15]). With a deep body and skull, weakly toothed mandibular jaws, and a vertical suspensorium, †*P. parvulus* diverges strongly from generalized actinopterygian conditions and likely had distinctive locomotor and feeding ecologies. Most significantly, we find that it possesses enlarged basibranchial tooth plates that oppose a tooth field contributed to by the median vomer and paired entopterygoids. Dating to ∼310 Ma, this represents the earliest candidate for a tongue-bite mechanism in actinopterygians, predating the oldest evidence for similar arrangements in extant groups by over 150 Myr. Modified gill-arch structure in †*Platysomus* amplifies the pattern of structural innovation among Carboniferous ray-fins, illustrates a step in the evolution of more elaborate tongue bites in related taxa, and further highlights the widespread convergence in actinopterygian feeding strategies across clades and over time.

## 2. Materials and Methods

### (a) Fossil specimens

†”Platysomidae” sensu Schultze et al.[16]

†*Platysomus superbus*. NHMUK PV P 4060: nearly complete articulated individual in part and counterpart, Glencartholm Volcanic Beds, Upper Border Group of the Calciferous Sandstone (Mississippian: Viséan), Glencartholm, Eskdale, Dumfriesshire, Scotland, UK.

†*Platysomus tenuistriatus*. NHMUK PV P 8029: impression of nearly complete articulated individual, Dalemoor Rake Ironstone, Pennine Lower Coal Measures Formation (Early Pennsylvanian: Bashkirian), Stanton-by-Dale, Staffordshire, England, UK.

†*Platysomus parvulus*. NHMUK PV P 11697: three-dimensionally preserved head and trunk broken in multiple pieces, Knowles Ironstone, Pennine Middle Coal Measures Formation (Early Pennsylvanian: Moscovian), Fenton, Staffordshire, England, UK (figures 1–2; supplementary material, figure 1).

**Fig. 1.**
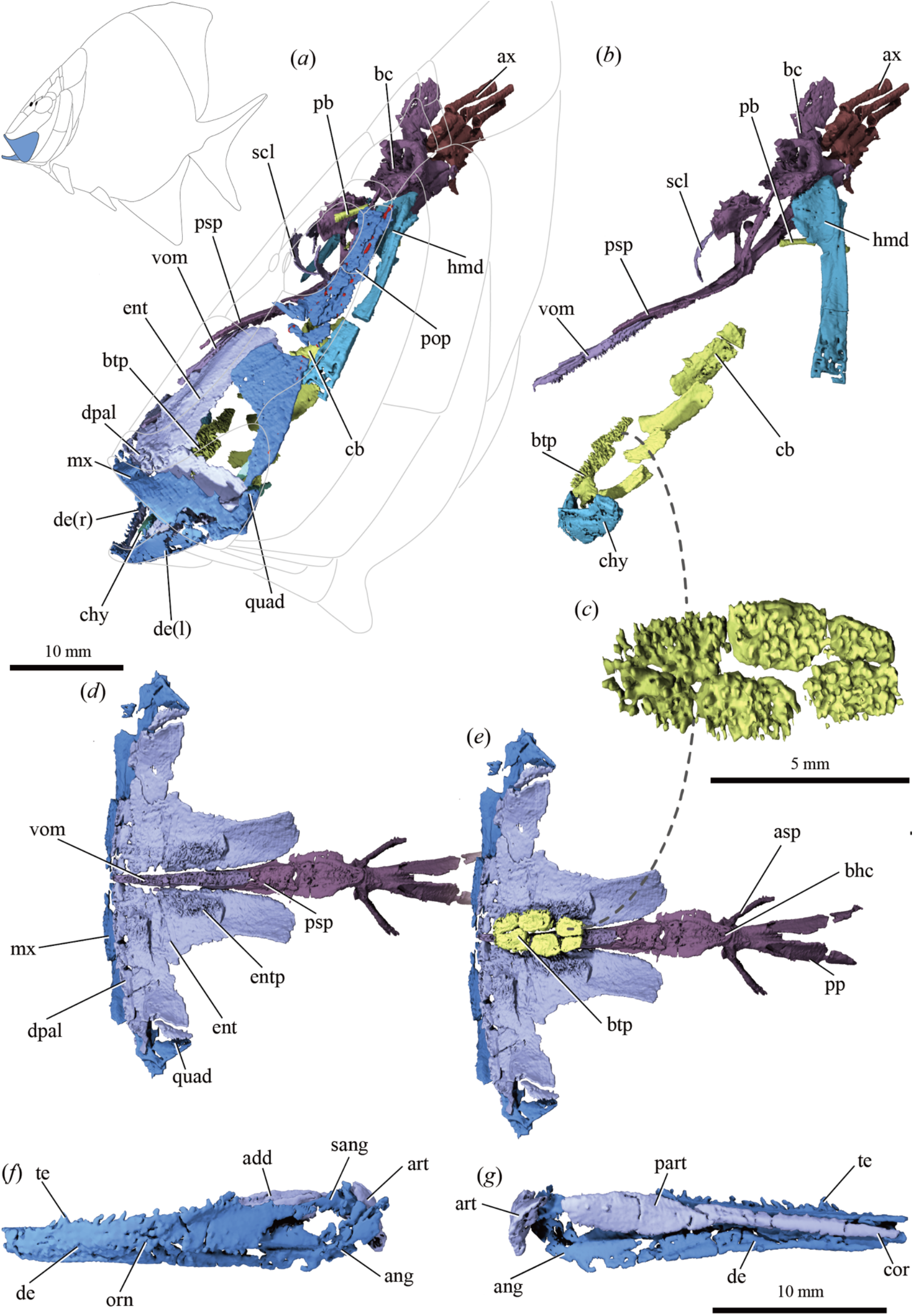
Cranial anatomy of †*Platysomus superbus* (NHMUK PV P11697) based on μCT scanning. (a) Bones of skull as preserved, in left lateral view; grey lines indicate external skull bones. Inset shows reconstruction of body shape, with maxilla and dentary in blue. (b) As in (a), but with dermal and jaw bones removed and elements repositioned. (c) Basibranchial tooth plate in dorsal view. (d) Upper jaw bones and parasphenoid in ventral view, with maxilla and palatal bones mirrored. (e) As in (d), but with basibranchial tooth plate shown in articulation. (f) Left mandible in lateral view and (g) medial view. Abbreviations: add, adductor fossa; ang, angular; art, articular; asp, ascending process of parasphenoid; ax, axial skeleton; bc, braincase; bhc, buccohypophysial canal; btp, basibranchial toothplate; cb, ceratobranchial; chy, ceratohyal; cor, coronoids; de, dentary; dpal, dermopalatine; ent, entopterygoid; entp, entopterygoid teeth; hmd, hyomandibula; mx, maxilla; orn, ornament; part, prearticular; pb, pharyngobranchial; pop, preoperculum; pp, posterior process of parasphenoid; psp, parasphenoid; quad, quadrate; sang, surangular; scl, sclerotic ossicle; te, teeth; vom, vomer. Colour coding: blue, cheek and outer jaw bones; light purple, palate and inner jaw bones; dark purple, braincase; turquoise, hyoid arch; yellow, branchial skeleton; brown, axial skeleton; red, lateral-line sensory canal pores.

†Bobasatraniidae sensu Schultze et al.[16]

†”*Platysomus*” *schultzei*. KUVP 86168: nearly complete individual in part and counterpart, Tinajas Member of the Atrasado Formation (Late Pennsylvanian: Kasimovian), Kinney Brick

Quarry, New Mexico, USA. The parasphenoid and upper and lower tooth plates are partially exposed on the surface of the part and counterpart. The left ascending process of the parasphenoid is broken and rotated at its base such that it lies parallel to the right ascending process (figure 2; supplementary material, figure 1).

**Fig. 2.**
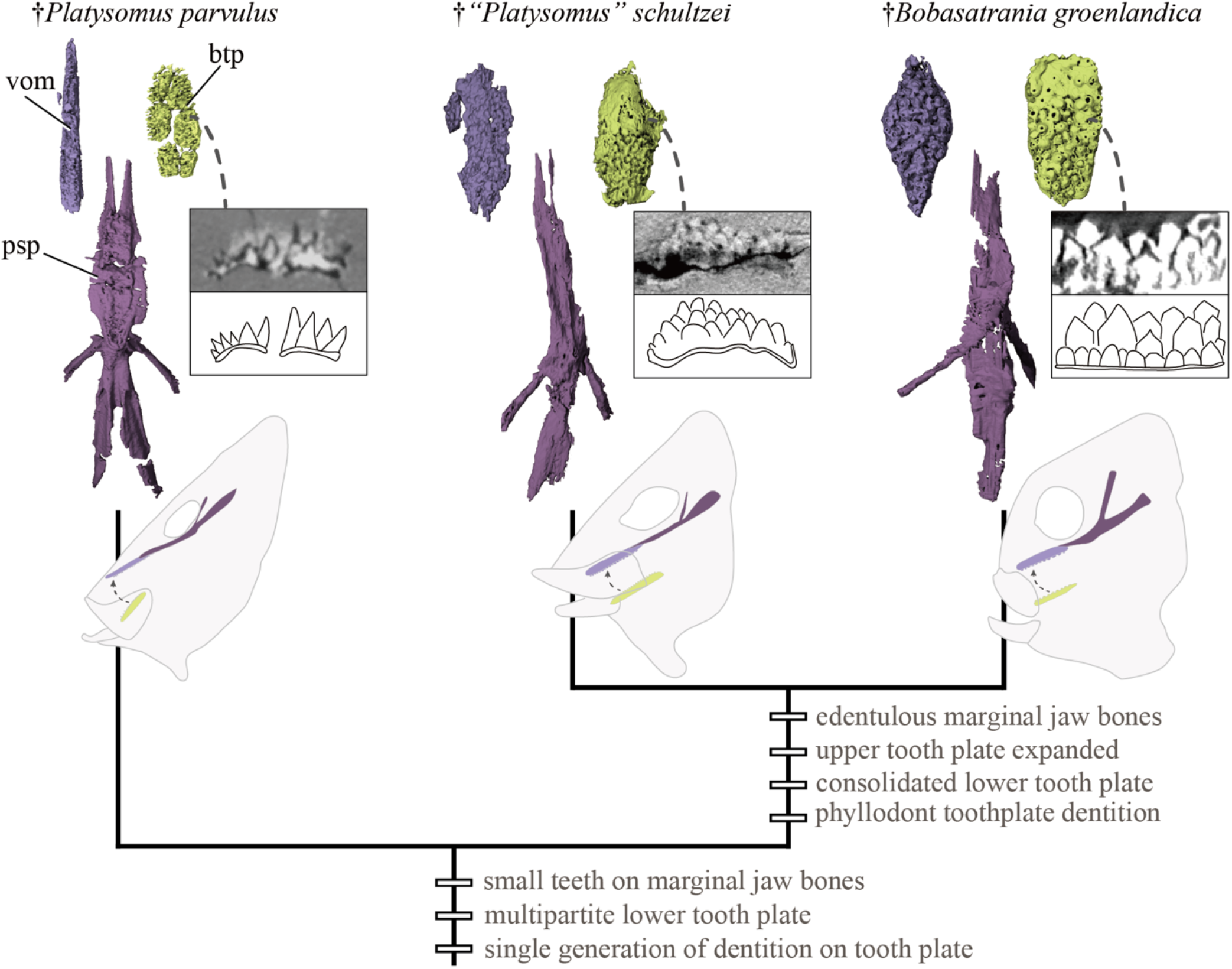
Evolution of a tongue bite in a lineage of Paleozoic to Mesozoic actinopterygians. Upper and lower tongue bite components in, from left to right, †*Platysomus superbus* (NHMUK PV P11697), † “*Platysomus*” *schultzei* (KUVP 86168), and †*Bobasatrania groenlandica* (NHMD 161449a,b). Upper row of insets show tomogram sections (upper) and schematic drawings (lower) through the lower tooth plate. Lower row of insets show schematic drawing of cranium and tongue bite components in lateral view. Abbreviations: btp, basibranchial tooth plate; psp, parasphenoid; vom, vomer. Colour coding: light purple, upper tooth plate; dark purple, parasphenoid; yellow, branchial tooth plate.

†*Bobasatrania groenlandica*. NHMD 161449a, b, nearly complete small individual in part and counterpart (figure 2; supplementary material, figure 2). Specimen is heavily pyritized in some regions, and pyrite has invaded the upper and lower tooth plates, obscuring the lower portions of some phyllodont tooth cusps. NHMD 303527a, skull portion of a large individual. Both specimens from the Wordie Creek Formation (Early Triassic: Induan), East Greenland.

†*Bobasatrania mahavavica*. NMS G.1956.17.15, NMS G.1956.17.16: nearly complete individual in part and counterpart from the Sakamena Group (Early Triassic: Induan), northwestern Madagascar.

### (b) Specimen visualization

We used Nikon XT H 225 ST micro-computed tomography scanners at the Imaging and Analysis Centre, Natural History Museum, London (NHMUK PV P 11697), CoLES CT Scanning Facility, University of Birmingham (NHMD 161449a, b and NHMD 303527a) and CTEES Facility, Department of Earth and Environmental Sciences, University of Michigan (KUVP 86168) to visualize internal structure in specimens. Scan parameters are provided in Supplementary Data. Materialise Mimics v.25 (Materialise Software, Leuven, Belgium; https://www.materialise.com/en/healthcare/mimics-innovation-suite/mimics), was used to segment these data, with exported surface meshes rendered in Blender v.2.79 (Blender Project; https://www.blender.org/). Some manual rearticulation of elements was carried out in Blender.

Mandibular, hyoid, and gill arch elements were realigned in †*Platysomus parvulus* (NHMUK PV P 11697) to account for postmortem splaying of these elements and a narrow crack running across the specimen. The parasphenoid and tooth plate elements in †”*Platysomus*” *schultzei* (KUVP 86168) are split across part and counterpart, and these were realigned; the distorted left ascending process was also restored to life position. Each slab consisted of multiple smaller blocks glued to one another; these were disarticulated and scanned separately for improved X-ray transmission.

## 3. Anatomical description

The specimen of †*Platysomus parvulus* (NHMUK PV P 11697) consists of a skull and anterior trunk preserved in multiple pieces. Slight anteroposterior compression has flared the upper and lower jaws, making the skull appear broader than in life. Consequently, the branchial apparatus is not crushed as in a laterally compressed individual. Cracks or missing fragments mean that some bones are only preserved on one side. We focus on the feeding apparatus (figures 1, 2; supplementary material, figure 3), with other internal details intended for a subsequent study.

**Fig. 3.**
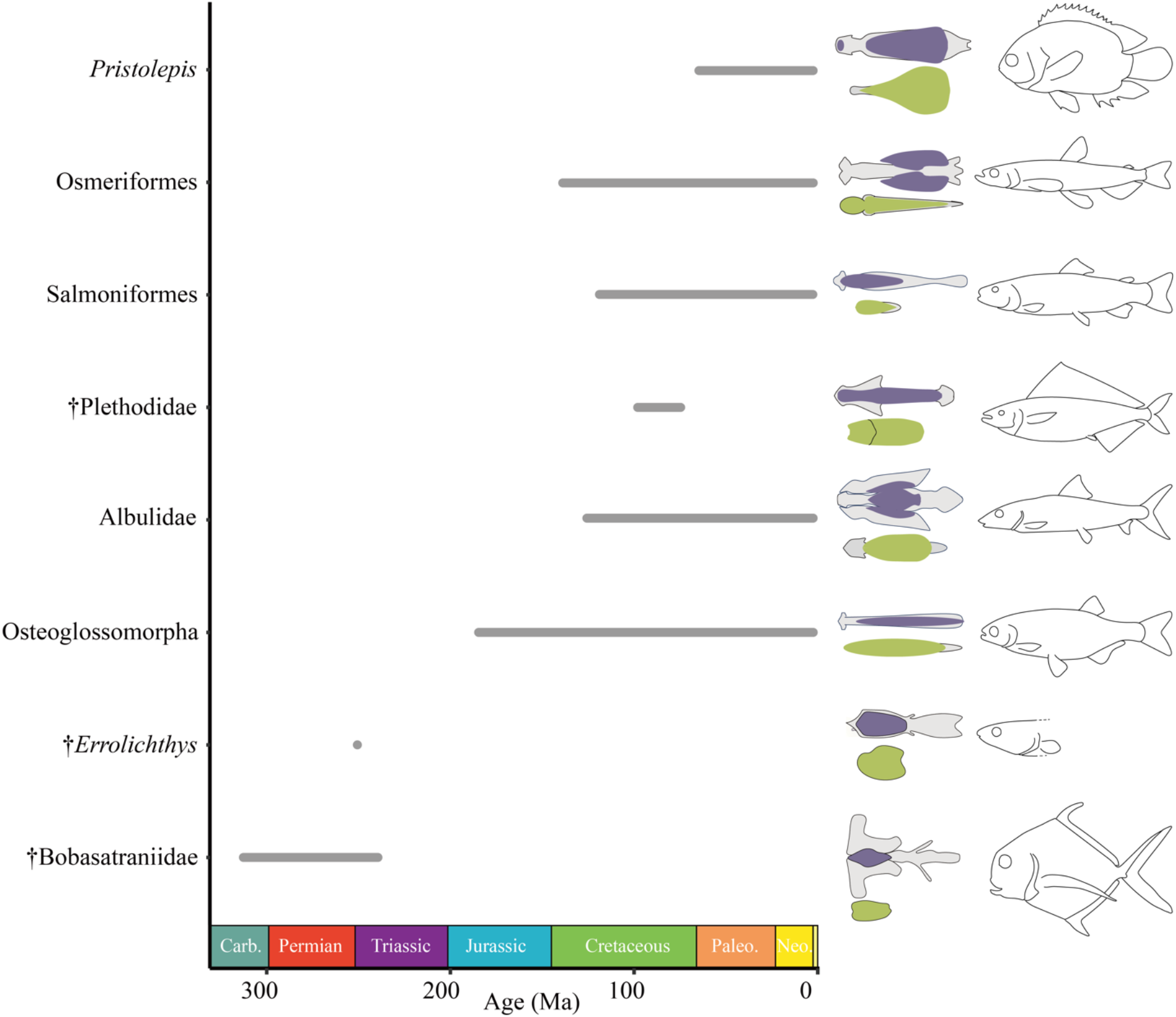
Multiple origins of tongue bite mechanisms in actinopterygians. Stratigraphic range (left), schematic drawing of tongue bite components (middle) and body profile (right) of actinopterygian lineages with tongue bites. Ranges for extinct taxa are based on fossil occurrences; ranges for extant taxa are based on molecular clock estimates of total-group ages from [49, 50]. Colour coding: grey, midline supporting skeletal elements; purple, upper dentigerous region; yellow, lower dentigerous region. Tongue bite schematics based on CT data (this paper; supplementary material, table 1) and [29-31]. Body profile schematics of extant taxa adapted from [51]; of extinct taxa adapted from [18, 30, 31, 52].

### (a) Palate including dorsal tooth plate and parasphenoid

A large “L”-shaped entopterygoid represents the principal bone of the palate. Its longer posterodorsal limb traces the profile of the parasphenoid and bears a broad, raised longitudinal band of teeth along its inner face. Four small dentigerous dermopalatines articulate in a row along the anteroventral margin of the entopterygoid. The trapezoidal ectopterygoid embraces the posteroventral limb of the entopterygoid, with a modest ectopterygoid process marking the anterior margin of the adductor chamber. The metapterygoid is unmineralized, with a small fragment of bone possibly representing a portion of the quadrate.

The parasphenoid and vomer extend along the palatal midline. Rather than being horizontal, they are oriented at a steep angle that approximates the profile of the skull. The spear-shaped median vomer is covered with a field of small, pointed teeth. When in articulation with the palate, these vomerine teeth form a continuous dental field with the longitudinal tooth bands of the entopterygoids. Tomograms show a single generation of teeth rather than superposed layers of older dentition (figure 2). The vomer inserts into a deep notch in the anterior end of the parasphenoid, posterior to which the latter broadens before sharply tapering anterior to the ascending processes. Small teeth, interrupted by a buccohypophysial opening, form a continuous band of dentition between this constriction and the vomer. There is no dermal basipterygoid process, but well-developed ascending processes sweep sharply posterolaterally and extend to the skull roof. They are tube-like and completely enclose the spiracular canal along their entire length. A long posterior stalk embraces the otic and occipital regions of the braincase, broadening as it approaches the occiput and split by a deep midline notch.

### (b) Hyoid arch, branchial skeleton, and ventral tooth plates

The hyoid arch includes a hyomandibula, single ceratohyal, and hypohyal. A small nodular endosketal bone near the jaw joint could represent an interhyal. The hyomandibula has a broad dorsal head and a long ventral shaft that is oriented vertically. Just ventral to the head, a canal pierces the shaft of the hyomandibula. A broad gap separates the distal tip of the hyomandibula from the single mineralized ceratohyal, which is short and plate like. The hypohyal is robust and gently curved in dorsal view.

Extensive mineralization appears limited to the ventral gill skeleton, which is most completely preserved on the left side. Four ossified ceratobranchials are present. The first three are slender and the fourth is substantially more robust. Three short, poorly preserved hypobranchials lie near the front of the gill skeleton. Median endoskeletal components are unmineralized. However, this region is covered by an oval dental field composed of six abutting tooth plates, arranged in two anteroposteriorly directed rows of three plates each (figure 1*c*). It appears that the individual plates are serially associated with arches one through three. This large composite plate opposes the dental field of the overlying vomer and entopterygoids and similarly bears a single generation of small, pointed teeth. Additional isolated tooth plates may be associated with other parts of the branchial skeleton. Mineralization of the dorsal skeleton is restricted to possible pharyngobranchials.

### (c) Mandible

The dentary constitutes most of the outer face of the slender mandible (figure 1*f, g*). It bears the mandibular sensory canal, prominent tuberculate external ornament, and a row of small conical teeth. An angular and possible surangular, plus a displaced articular, make up the posterior end of the jaw. The inner face of the mandible is lined with an indeterminate number of coronoids plus a lozenge-shaped prearticular that bears a lateral process defining the anterior margin of the adductor fossa. These inner bones bear very fine denticles that are beyond the limits of segmentation.

## 4. Discussion

### (a) Evolution of tongue-bite mechanisms in Paleozoic actinopterygians

Our three-dimensional tomographic data for †*Platysomus parvulus* provide the earliest evidence of a tongue bite in actinopterygians (figure 3). Other early examples come from younger bobasatraniids (figure 2): the Late Pennsylvanian (Kasimovian; ∼305 Ma)

†”*Platysomus*” *schultzei* bears an expanded vomerine tooth plate plus a single ventral one [17], with a similar, but more substantial, mechanism apparent in three-dimensional Early Triassic material of †*Bobasatrania mahavavica* and †*B. groenlandica* [18]. While *in situ* tooth plates are rare, abundant occurrences of isolated examples are known from the Late Pennsylvanian to the Middle Triassic [17, 19, 20].

The tooth plates of the older †*P. parvulus* correspond positionally to those of †bobasatraniids, but there are important structural differences (figure 2). First, the vomerine plate is narrow and relatively flat, rather than broad and convex in younger taxa. Second, the faintly convex basibranchial plate is composed of two rows of three separate but tightly adjoining plates, in contrast to a single massive plate in †bobasatraniids. Third and finally, both the upper and lower plates of †*P. parvulus* bear only a single layer of pointed cusps, in contrast to the multiple superimposed generations seen in bobasatraniids (so-called “phyllodont” plates [19, 21]). For all three features, the condition in †*P. parvulus* appears primitive relative to †bobasatraniids. Branchial tooth plates that are small, flat, and bear a single layer of teeth or denticles are widespread among early osteichthyans, with basibranchial tooth plates typically consisting of paired rather than median bones [22].

The plesiomorphic construction of dental plates of †*P. parvulus* suggests a role in understanding the evolution of the more elaborate tongue bites of †bobasatraniids. A close relationship between at least some †platysomids and †bobasatraniids has previously been proposed [9, 16, 17, 19, 23, 24]. We find additional anatomical support for this hypothesis not only from the tongue bite, but also by the highly unusual enclosure of the spiracular canal within the body of the ascending process of the parasphenoid. However, †*P. parvulus* lies outside of †bobasatraniids (sensu Schultze et al. [16]) based on features including the absence of a highly modified quadratojugal and the presence of long, tooth-bearing jaws (figure 1*f, g*) rather than short edentulous ones (supplementary material, figure 5). These primitive mandibular traits suggest that †*P. parvulus* retained use of the jaws in biting, with additional prey manipulation and reduction provided by the basibranchial dental plates. This stands in contrast to what seems to be the exclusive use of a tongue bite in †bobasatraniids, with the toothless jaws— which may have been kinetic in some species [18, 25]—instead dedicated to governing fluid flow into the oral chamber. It is possible that shifting food processing to the branchial skeleton lifted constraints on the mandibular jaws in later †bobasatraniids, allowing them to become more mobile. Therefore †*P. parvulus* captures a key intermediate stage for one of the most specialized feeding mechanisms to evolve among Paleozoic actinopterygians.

### (b) Tongue bites as a recurrent and versatile functional innovation

Tongue bite mechanisms have evolved numerous times in actinopterygian lineages, including in several entirely extinct groups [22], with substantial variation in the number of tooth-bearing palatal and branchial elements and the extent of the dentigerous covering (figure 3) [26-28]. Although principally applied to the mechanism in osteoglossomorphs, there has been widespread use of the term in the literature. Here we broadly define a tongue bite as the use of midline elements of the gill skeleton opposing toothed surfaces on the roof of the mouth. Other lineages, like galaxiiforms and argentiniforms, display prominent ventral midline dentition on the anterior gill elements, but appear to lack—or have lost—the dorsal dentigerous elements of a tongue bite [28]. The tongue bite seen in †*P. parvulus* and later †bobasatraniids is the earliest known example of this mechanism in actinopterygians, which has evolved numerous times, including in several entirely extinct groups. The enigmatic Early Triassic †*Errolichthys* represents the next oldest case of an actinopterygian tongue bite, with a large oval basihyal dental plate opposing a similar field on the parasphenoid [29, 30]. Late Cretaceous †plethodids represent the most diverse extinct group characterized by a tongue bite, with nearly 20 genera known from this early diverging crown teleost lineage [31]. These are joined by additional isolated examples in Mesozoic teleost groups (e.g., †*Cimolichthys*; [32]).

The tongue bite mechanism of living teleosts provides an actualistic model for interpreting function in †*P. parvulus* and †bobasatraniids. Osteoglossomorphs (bony tongues) are the most celebrated extant tongue-biting group, with many species bearing prominent fangs on the parasphenoid, basihyal, or both [3]. A tongue bite involving fangs is also present in salmoniforms, with *in vivo* kinematic studies of both groups showing prey reduction via “raking” [4, 33]. Tongue bites incorporating low crowns rather than long pointed teeth are better structural—and presumably functional—analogues for examples in Paleozoic actinopterygians. In *Pristolepis* (leaffish), the basihyal plate and parasphenoid have opposing fields of rounded teeth [34]. Along with additional adjacent pharyngeal tooth plates, these contribute to a complex chewing cycle [34] for breaking down the cuticle of insects and insect larvae [35]. *Albula* (bonefish) provides the best analogue for Paleozoic tongue biters. It possesses phyllodont basibranchial and parasphenoid plates with robust, rounded teeth superposed in several layers, with a rich fossil record of these phyllodont plates extending into the Cretaceous [21]. By weight,

*Albula* stomach contents are dominated by hard prey like crustaceans and mollusks [36]. This suggests durophagy in †bobasatraniids, further supported by wear on the dental plates of †*B. scutata* [20].

### (c) Tongue bites as a component of the early trophic radiation of actinopterygians

Actinopterygians show substantial expansion in taxonomic and morphological diversity in the Carboniferous relative to the Devonian [8, 37]. Past work suggests this shift might relate to diversification spurred by ecological opportunities opened by the Devonian-Carboniferous extinction [38, 39]. In particular, the Carboniferous records a proliferation of tooth and jaw traits bearing on feeding ecology. These include the appearance of coronoid processes (several taxa; [40, 41]); styliform (†Guildayichthyiformes [42]), peg-like (†*Frederichthys* [43]), and bulbous (†*Mesolepis* [44]) teeth, strongly heterodont dentitions (†*Paphosiscus* [45]), large fangs (several potentially unrelated lineages; [12]), and dense dental batteries, consolidated dental plates, and beaks (†Eurynotiformes; [9, 10]). Many represent the first examples of traits that would go on to appear independently in later, distantly related ray-fin lineages.

Tongue bites represent an additional feeding innovation in Carboniferous actinopterygians, but their first appearance postdates many other examples. The oldest †platysomids preserving cranial material date to the mid-Mississippian (mid Viséan; ∼338 Ma [46]) but show no evidence of opposing dorsal and ventral dental plates, while the arrangement in the younger †*P. parvulus* suggests it might not be long removed from the origin of the tongue bite itself. However, the multipartite construction of its primitive dorsal and ventral plates suggest that isolated components of early tongue bites might not be found or recognized, so we cannot exclude an earlier origin. This contrasts with the relatively continuous record of phyllodont plates of related †bobasatraniids in late Pennsylvanian to Middle Triassic deposits, representing a duration of ∼70 Myr for this successful lineage of hard-prey specialists. This delayed origin of a crushing tongue bite relative to the Devonian-Carboniferous transition contrasts with the pattern for the other principal lineage of durophagous Paleozoic actinopterygians: †eurynotiforms. The earliest †eurynotiforms already show specializations for consuming hard prey in the Tournaisian [9], and by the mid Viséan taxa like †*Cheirodopsis* possess highly consolidated upper and lower tooth plates and beak-like oral jaws [46]. The offset origins of these different anatomical solutions to the shared problem of processing hard prey reinforces other evidence for a protracted interval of innovation in early actinopterygians, spanning the Carboniferous and potentially extending into the latest Devonian [47, 48].

Ultimately, placing these functional traits within an explicitly phylogenetic context will be necessary for testing whether feeding innovations arose at a steady rate or instead show an evolutionary burst consistent with ecological release.

## Acknowledgements

We thank B. Lindow (NHMD), C. Beard and M. Sims (KUVP), E. Bernard and Z. Johanson (NHMUK), and S. Walsh (NMS) for access to specimens. We thank V. Fernandez and B. Clark (Imaging and Analysis Centre, NHMUK) for assistance with CT scanning. W. Sanders (University of Michigan Museum of Paleontology) disarticulated and re-articulated KUVP 86168.

SG was supported by a Royal Society Dorothy Hodgkin Research Fellowship (no. DH160098) and the National Environment Research Council (NE/X016633/1). MF was supported by the National Science Foundation (EAR 2219007 and DEB 2333684). MAK was supported by the National Science Foundation (DEB 2333683).

## Data availability

Reconstructed tomogram stacks (as .TIFF files) and surface meshes for 3D segmented elements (as .PLY and.OBJ files) of specimens scanned for this study are archived on MorphoSource and Zenodo; URL links are provided during the review process and will be replaced with stable Morphosource DOIs upon acceptance. Full details are given in the Supplementary Information.

## Supplementary Information

### Computed tomography parameters

†*Platysomus parvulus* NHMUK PV P 11697 was scanned using a Nikon XT H 225 at the Imaging and Analysis Centre, Natural History Museum, London, with the following settings: reflection target, voltage, 200 kV; current, 180 μA; exposure, 1 s; projections, 3141; frames per projection, 4; filter, 0.5 mm copper; effective voxel size, 31.03 μm; option for minimizing ring artefacts was selected.

†”*Platysomus*” *schultzei* (KUVP 86168) was scanned twice using a Nikon XT H 225 at the CTEES Facility, Department of Earth and Environmental Sciences, University of Michigan. The first scan included both the part and counterpart and captured the entire cranial region, with the following settings: voltage, 197 kV; current, 180 μA; exposure, 4 s; projections, 3141; frames per projection, 2; filter, 3.25 mm copper; effective voxel size, 40.86 μm; option for minimizing ring artefacts was selected. The second scan focussed on the left lower jaw preserved in the part, with the following settings: voltage, 120 kV; current, 102 μA; exposure, 1 s; projections, 3141; frames per projection, 2; filter, none; effective voxel size, 12.32 μm.

†*Bobasatrania groenlandica* NHMD 161449a, b was scanned with part and counterpart together using a Nikon XT H 225 ST 2x at the CoLES CT Scanning Facility, University of Birmingham, using the following settings: rotating target, voltage, 220 kV; current, 229 μA; exposure, 1 s; projections, 4492; frames per projection, 2; filter, 1 mm tin; effective voxel size, 16.89 μm.

## Data availability

CT data and segmented 3D objects for the studied specimens are available on Morphosource (stable DOIs will be generated upon acceptance):

†*Platysomus parvulus*NHMUK PV P 11697

Reconstructed TIFF stack:

https://www.morphosource.org/concern/media/000740374?locale=en

OBJ file of all segmented three-dimensional objects, which can be manipulated as individual PLY files when opened in Blender:

https://www.morphosource.org/concern/media/000740377?locale=en

†”*Platysomus*” *schultzei*KUVP 86168

Reconstructed TIFF stack of head region:

https://www.morphosource.org/concern/media/000740358?locale=en

OBJ file of all segmented three-dimensional objects in the head region, which can be manipulated as individual PLY files when opened in Blender:

https://www.morphosource.org/concern/media/000740361?locale=en

Reconstructed TIFF stack of jaw region:

https://www.morphosource.org/concern/media/000740338?locale=en

PLY file of segmented jaw:

https://www.morphosource.org/concern/media/000740364?locale=en

†*Bobasatrania groenlandica*NHMD 161449a, b

Reconstructed TIFF stack:

https://www.morphosource.org/concern/media/000740354?locale=en

OBJ file of all segmented three-dimensional objects, which can be manipulated as individual PLY files when opened in Blender:

https://www.morphosource.org/concern/media/000740367?locale=en

Mimics files for the above are available on Zenodo (a stable DOI will be generated upon acceptance): https://zenodo.org/records/15380086?preview=1&token=eyJhbGciOiJIUzUxMiJ9.eyJpZCI6IjhjOTkwY2E2LTBjNWQtNDRkNy04MDQ5LThlMDI1Mzg1ODAwMyIsImRhdGEiOnt9LCJyYW5kb20iOiJkOGM3ZGY2N2Y4MzA4NGNiZDJmMWYyNmViOGM4YjJjOSJ9.H7aPEkOwa1PQTQI9xeyfBd5umLfFiP1Ip4q11w7Scejfpxf6mKdUmr79I5XLTfcMz1obJc780ypBMEymLeZrTA

## Supplementary figures

**S Fig. 1.**
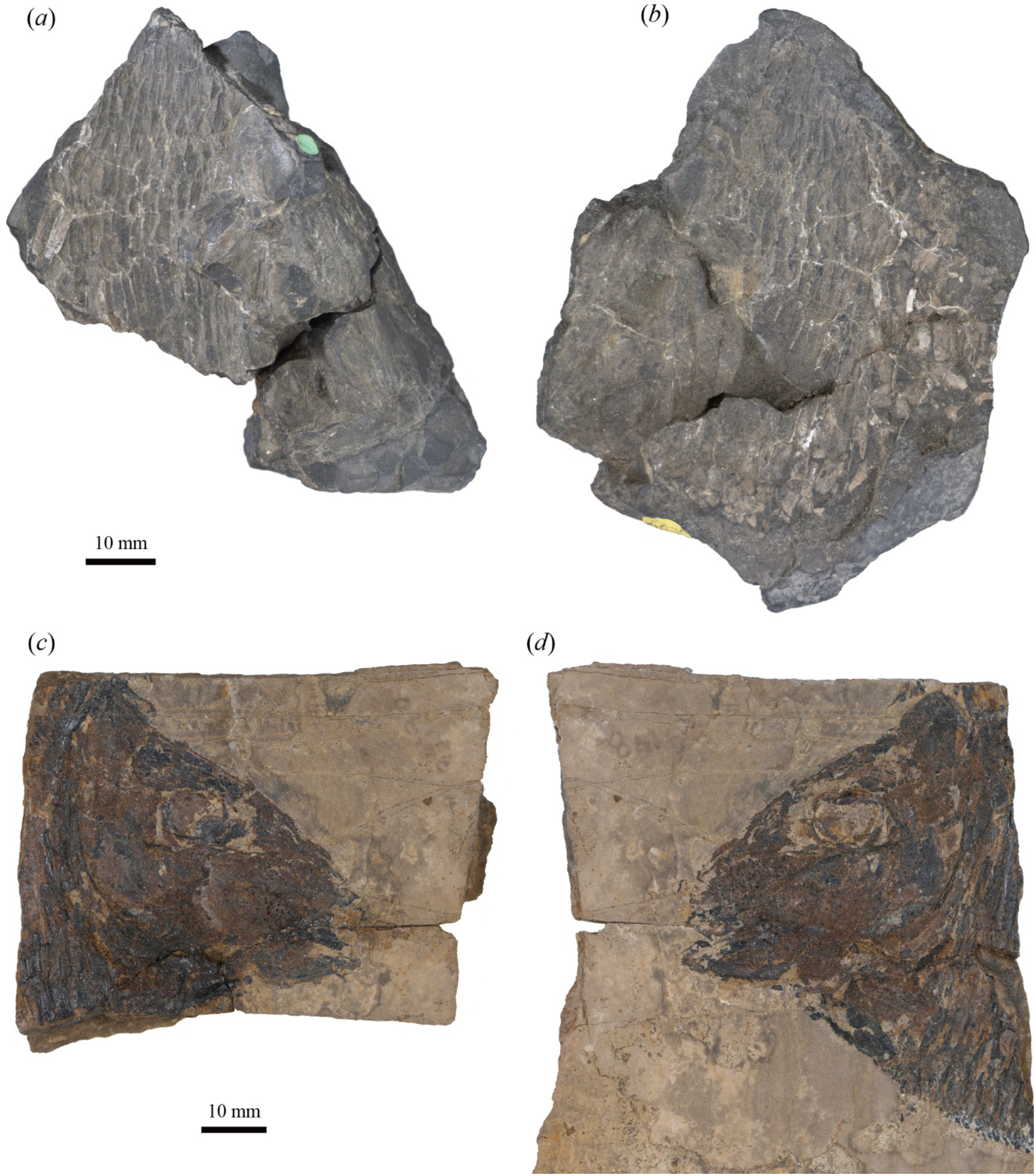
External photographs of specimens. †*Platysomus parvulus* (NHMUK PV P11697) in (a) part and (b) counterpart; †”*Platysomus*” *schultzei* (KUVP 86168) in (c) part and (d) counterpart.

**S Fig. 2.**
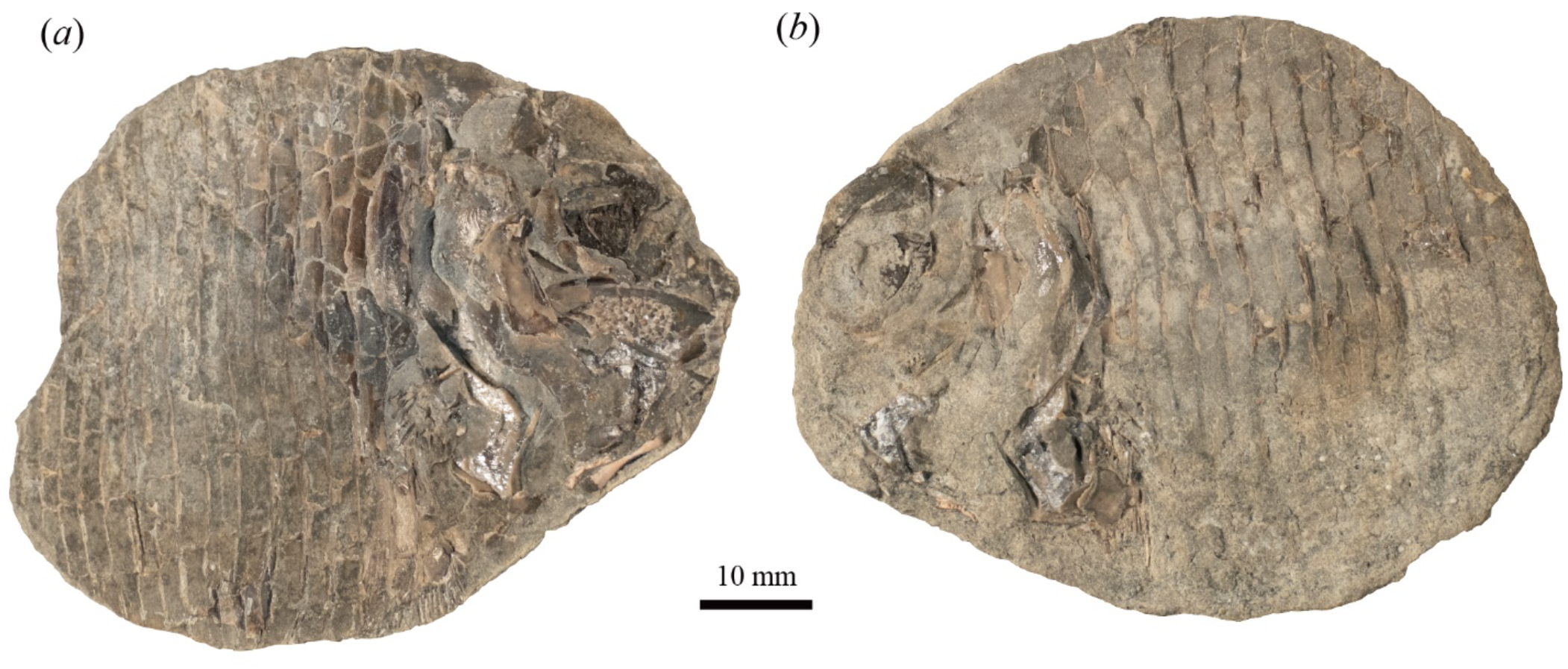
External photographs of specimens. †*Bobasatrania groenlandica* (NHMD 161449a, b) in (a) part and (b) counterpart.

**S Fig. 3.**
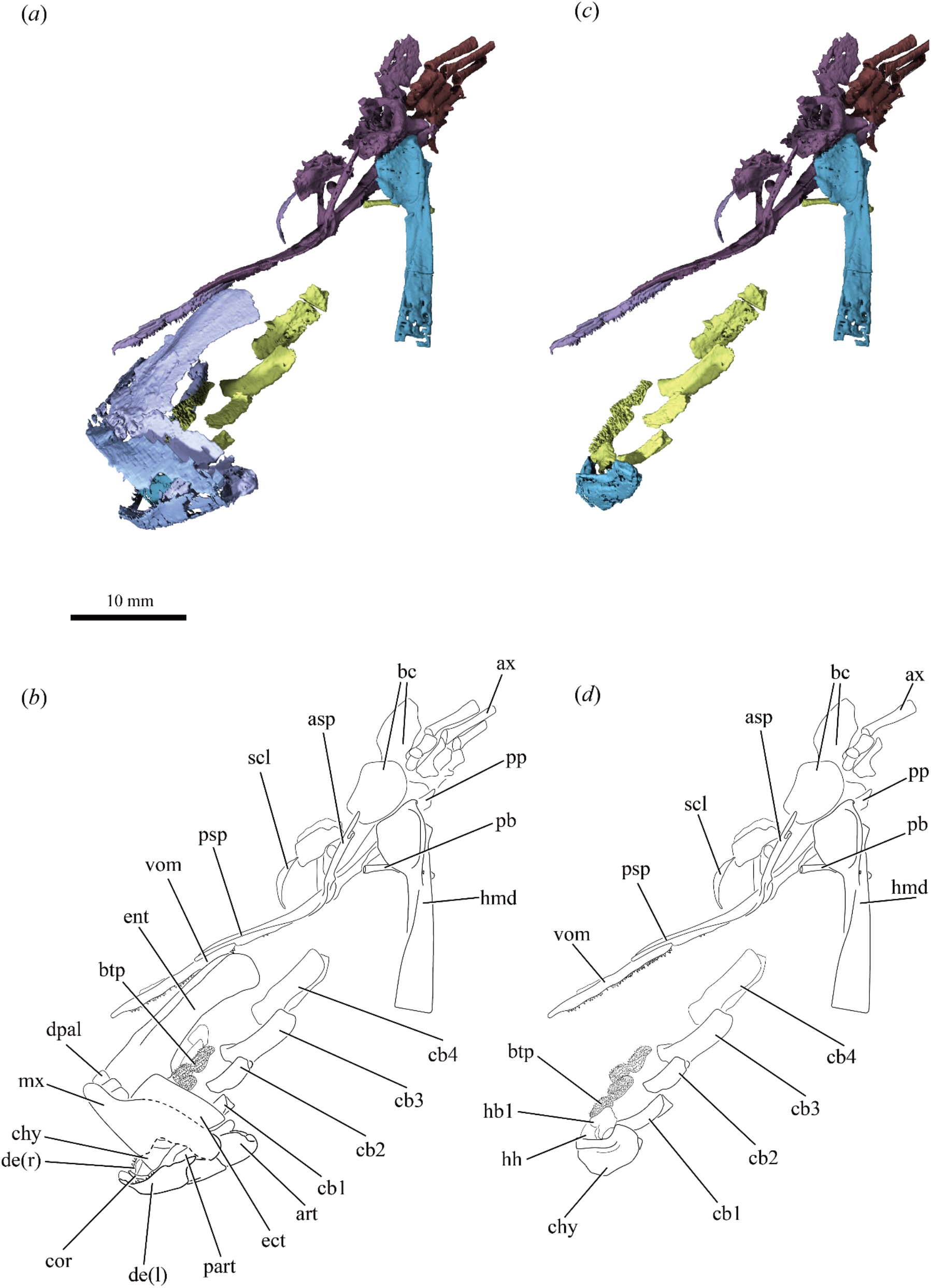
Cranial anatomy of †*Platysomus parvulus* (NHMUK PV P11697) based on μCT scanning. Render (a) and interpretive drawing (b) of repositioned bones of skull, with dermal cheek elements removed. Render (c) and interpretive drawing (d) of repositioned bones of skull, with dermal cheek and jaw elements removed. Abbreviations: art, articular; asp, ascending process of parasphenoid; ax, axial skeleton; bc, braincase; btp, basibranchial toothplate; cb, ceratobranchial; chy, ceratohyal; cor, coronoids; de, dentary; dpal, dermopalatine; ent, entopterygoid; hb, hypobranchial; hh, hypohyal; hmd, hyomandibula; mx, maxilla; part, prearticular; pb, pharyngobranchial; pp, posterior process of parasphenoid; psp, parasphenoid; scl, sclerotic ossicle; vom, vomer. Colour coding: blue, cheek and outer jaw bones; light purple, palate and inner jaw bones; dark purple, braincase; turquoise, hyoid arch; yellow, branchial skeleton; brown, axial skeleton; red, lateral-line sensory canal pores.

**S Fig. 4.**
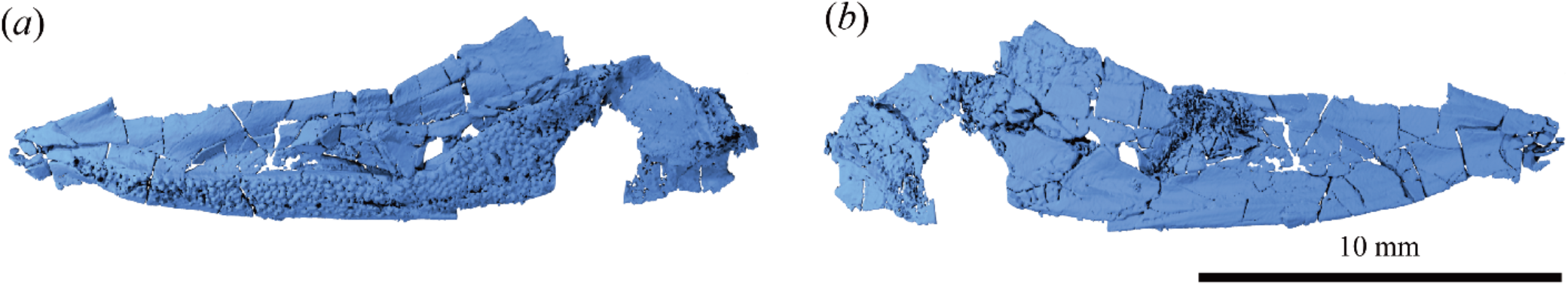
Left lower jaw of †*”Platysomus” schultzei* (KUVP 86168). (a) lateral view. (b) medial view.

## Supplementary tables

**S Table 1.** Skeletal elements involved in the tongue bite apparatuses of extant fishes.

| Genus | Family | Clade | Basihyal | Basibranchial | Other | Parasphenoid | Ectopterygoid | Other | or Specimen |
| --- | --- | --- | --- | --- | --- | --- | --- | --- | --- |
| *Albula* | Albulidae | Elopomorpha | X | X | - | X | X | - | UF 207902, MorphoSource |
| *Spirinchus* | Osmeridae | Protacanthopterygii | X | X | - | 0 | X | - | UW 166548, MorphoSource |
| *Pristolepis* | Pristolepididae | Percomorpha | X | X | 2nd Cb | X | 0 | Vomer | USNM 389747, UMMZ Scan |
| *Hiodon* | Hiodontidae | Osteoglossomorpha | X | X | - | X | X | Palatine + Entopterygoid | Hilton, 2001 |
| *Oncorhynchus* | Salmonidae | Salmoniformes | X | X | 2nd Cb | X | X | - | UW 018741, Morphosource |

## References

[1] Wainwright, P.C. 2005 Functional morphology of the pharyngeal jaw apparatus. Fish Physiology 23, 77–101.

[2] Mehta, R.S. & Wainwright, P.C. 2007 Raptorial jaws in the throat help moray eels swallow large prey. Nature 449, 79–82.

[3] Hilton, E.J. 2001 Tongue bite apparatus of osteoglossomorph fishes: variation of a character complex. Copeia 2001, 372–381.

[4] Camp, A.L., Konow, N. & Sanford, C.P.J. 2009 Functional morphology and biomechanics of the tongue-bite apparatus in salmonid and osteoglossomorph fishes. Journal of Anatomy 214, 717–728.

[5] Liem, K.F. 1973 Evolutionary strategies and morphological innovations: cichlid pharyngeal jaws. Systematic Zoology 22, 425–441.

[6] Friedman, M. 2022 The macroevolutionary history of bony fishes: a paleontological view. Annual Review of Ecology, Evolution, and Systematics 53, 353–377.

[7] Vermeij, G.J. 1977 The Mesozoic marine revolution: evidence from snails, predators and grazers. Paleobiology 3, 245–258.

[8] Sallan, L.C. & Friedman, M. 2012 Heads or tails: staged diversification in vertebrate evolutionary radiations. Proceedings of the Royal Society B 279, 2025–2032.

[9] Sallan, L.C. & Coates, M.I. 2013 Styracopterid (Actinopterygii) ontogeny and the multiple origins of post-Hangenberg deep-bodied fishes. Zoological Journal of the Linnean Society 169, 156–199.

[10] Friedman, M., Pierce, S.E., Coates, M.I. & Giles, S. 2019 Feeding structures in the ray-finned fish Eurynotus crenatus (Actinopterygii: Eurynotiformes): implications for trophic diversification among Carboniferous actinopterygians. Earth and Environmental Science Transactions of the Royal Society of Edinburgh 109, 33–47.

[11] Gardiner, B.G. 1984 The relationships of the palaeoniscid fishes, a review based on new species of Mimia and Moythomasia from the Upper Devonian of Western Australia. Bulletin of the British Museum (Natural History) Geology 37, 173–428.

[12] Friedman, M., Figueroa, R.T., Hodnett, J.-P., Lucas, S.G., Higgins, R.R., Pierce, S. & Giles, S. 2024 A new genus and species of large macrodont actinopterygian from the Pennsylvanian (Kasimovian/Missourian) Atrasado Formation of New Mexico. Contributions from the Museum of Paleontology, University of Michigan 36, 8–42.

[13] Watson, D.M.S. 1925 The structure of certain palæoniscids and the relationships of that group with other bony fishes. Proceedings of the Zoological Society of London 54, 815–870.

[14] Giles, S., Darras, L., Clément, G., Blieck, A. & Friedman, M. 2015 An exceptionally preserved Late Devonian actinopterygian provides a new model for primitive cranial anatomy in ray-finned fishes. Proceedings of the Royal Society B 282, 2015.1485.

[15] Patterson, C. & Rosen, D.E. 1977 A review of ichthyodectiform and other Mesozoic teleost fishes and the theory and pratice of classifying fossil. Bulletin of the American Museum of Natural History 158, 81–172.

[16] Schultze, H.-P., Mickle, K.E., Poplin, C., Hilton, E.J. & Grande, L. 2021 Actinopterygii I: Palaeoniscimorpha, Stem Neopterygii, Chondrostei. Munich, Verlag Dr. Friedrich Pfeil; 299 p.

[17] Zidek, J. 1992 Late Pennsylvanian Chondrichthyes, Acanthodii, and deep-bodied Actinopterygii from the Kinney Quarry, Manzanita Mountains, New Mexico. In Geology and paleontology of the Kinney Brick Quarry, Late Pennsylvanian, central New Mexico (ed. J. Zidek), pp. 145–182.

[18] Nielsen, E. 1952 A preliminary note on Bobasatrania groenlandica. Meddelelser fra Dansk Geologisk Forening 12, 197–204.

[19] Johnson, G.D. & Zidek, J. 1981 Late Paleozoic phyllodont tooth plates. Journal of Paleontology 55, 524–536.

[20] Böttcher, R. 2014 Phyllodont tooth plates of Bobasatrania scutata (Gervais, 1852) (Actinoperygii, Bobasatraniiformes) from the Middle Triassic (Longobardian) Grenzbonebed of southern Germany and eastern France, with an overview of Triassic and Palaeozoic phyllodont tooth plates. Neues Jarhbuch für Geologie und Paläontologie, Abhandlungen 274, 291–311.

[21] Estes, R. 1969 Studies on fossil phyllodont fishes: interrelationships and evolution in the Phyllodontidae (Albuloidei). Copeia 1969, 317–331.

[22] Nelson, G.J. 1969 Gill arches and the phylogeny of fishes: with notes on the classification of vertebrates. Bulletin of the American Museum of Natural History 141, 475–552.

[23] Wilson, C.D., Mansky, C.F. & Anderson, J.S. 2021 A platysomid occurrence from the Tournaisian of Nova Scotia. Scientific Reports 11, 8375.

[24] Campbell, K.S.W. & Phuoc, L.D. 1983 A Late Permian actinopterygian fish from Australia. Palaeontology 26, 33–70.

[25] Tintori, A., Hitij, T., Jiang, D., Lombardo, C. & Sun, Z. 2014 Triassic actinopterygian fishes: the recovery after the end-Permian crisis. Integrative Zoology 9, 394–411.

[26] Gosline, W.A. 1971 Functional Morphology and Classification of Teleostean Fishes. Honolulu, University of Hawaii Press; 208 p.

[27] Weitzman, S.H. 1967 The origin of the stomiatoid fishes with comments on the classification of salmoniform fishes. Copeia 1967, 507–540.

[28] Williams, L.R.G. 1997 Bones and muscles of the suspensorium in the galaxioids and Lepidogalaxias salamandroides (Teleostei: Osmeriformes) and their phylogenetic significance. Records of the Australian Museum 42, 139–166.

[29] Lehman, J.-P. 1952 Étude complémentaire des Poissons de l’Eotrias de Madagascar. Kungliga Svenska Vetenskaps-Akademiens Handlingar, Ser. IV 2, 1–201.

[30] Nielsen, E. 1955 Notes on Triassic fishes from Greenland. I. Errolichthys mirablis Lehman. Meddelelser fra Dansk Geologisk Forening 11, 563–578.

[31] Taverne, L. & Gayet, M. 2004 Ostéologie et relations phylogénétiques des Protobramidae (Teleostei, Tselfatiiformes) du Cénomanien (Crétacé supérieur) du Liban. Cybium 28, 285–314.

[32] Beckett, H.T., Giles, S.G. & Friedman, M. 2018 Comparative anatomy of the gill skeleton of fossil Aulopiformes (Teleostei: Eurypterygii). Journal of Systematic Palaeontology 16, 1221–1245.

[33] Konow, N. & Sanford, C.P.J. 2008 Biomechanics of a convergently derived prey-processing mechanism in fishes: evidence from comparative tongue bite apparatus morphology and raking kinematics. Journal of Experimental Biology 211, 3378–3391.

[34] Liem, K. & Greenwood, P.H. 1983 A functional approach to the phylogeny of pharyngognath teleosts. American Zoologist 21, 83–101.

[35] Sangpradub, N. & Hanjavanit, C. 2017 Diet Composition of Pristolepis fasciata (Bleeker, 1851) (Family Nandidae) and Puntius brevis (Bleeker, 1849) (Family Cyprinidae) in Kaeng Lawa, Thailand. Chiang Mai Journal of Science 44, 839–846.

[36] Crabtree, R.E., Stevens, C., Snodgrass, D. & Stengard, F.J. 1998 Feeding habits of bonefish, Albula vulpes, from the waters of the Florida Keys. Fishery Bulletin 96, 754–766.

[37] Henderson, S., Dunne, E.M., Fasey, S.A. & Giles, S. 2023 The early diversification of ray-finned fishes (Actinopterygii): hypotheses, challenges, and future prospects. Biological Reviews 98, 284–315. (doi:10.31223/X58D1D).

[38] Sallan, L.C. & Coates, M.I. 2010 End-Devonian extinction and a bottleneck in the early evolution of modern jawed vertebrates. Proceedings of the National Academy of Sciences of the USA 107, 10131–10135.

[39] Friedman, M. & Sallan, L.C. 2012 Five hundred million years of extinction and recovery: a Phanerozoic survey of large-scale diversity patterns in fishes. Palaeontology 55, 707–742.

[40] Gottfried, M.D. 1992 Functional morphology of the feeding mechanism in a primitive paleonsicoid-grade actinopterygian fish. In Fossil Fishes as Living Animals (ed. E. Mark-Kurik), pp. 151–158. Tallinn, Estonia, Academy of Sciences.

[41] Coates, M.I. & Tietjen, K. 2019 ‘This strange little palaeoniscid’: a new early actinopterygian genus, and commentary on pectoral fin conditions and function. Earth and Environmental Science Transactions of the Royal Society of Edinburgh 109, 15–31.

[42] Lund, R. 2000 The new actinopterygian order Guildayichthyiformes from the Lower Carboniferous of Montana (USA). Geodiversitas 22, 171–206.

[43] Coates, M.I. 1998 Actinoptergians from the Namurian of Bearsden, Scotland, with comments on early actinopterygian neurocrania. Zoological Journal of the Linnean Society 122, 27–59.

[44] Elliott, F.M. & Giles, S. In press A new species of Mesolepis (Actinopterygii) from the Late Carboniferous of Scotland, with especial reference to Mesolepis wardi Young. Earth and Environmental Science Transactions of the Royal Society of Edinburgh.

[45] Grogan, E. & Lund, R. 2015 Two new Actinopterygii (Vertebrata, Osteichthyes) with cosmine from the Bear Gulch Limestone (Heath Fm., Serpukhovian, Mississippian) of Montana USA. Proceedings of the Academy of Natural Sciences of Philadelphia 164, 111–132.

[46] Moy-Thomas, J.A. & Bradley-Dyne, M. 1938 The actinopterygian fishes from the Lower Carboniferous of Glencartholm, Eskdale, Dumfriesshire. Transactions of the Royal Society of Edinburgh 59, 437–480.

[47] Giles, S., Feilich, K.L., Warnock, R.C.M., Pierce, S.E. & Friedman, M. 2023 A Late Devonian actinopterygian suggests high lineage survivorship across the end-Devonian mass extinction. Nature Ecology & Evolution 7, 10–19.

[48] Igielman, B., Figueroa, R.T., Higgins, R.R., Pierce, S., Coates, M.I., Troyer, E.M., Fernandez, V., Dollman, K., Lu, J., Zhu, M., et al. In review The lower jaw of Devonian ray-finned fishes (Actinopterygii): anatomy, relationships, and functional morphology.

[49] Rabosky, D.L., Chang, J., Title, P.O., Cowman, P.F., Sallan, L., Friedman, M., Kaschner, K., Garilao, C., Near, T.J., Coll, M., et al. 2018 An inverse latitudinal gradient in speciation rate for marine fishes. Nature 559, 392–395.

[50] Harrington, R.C., Kolmann, M.A., Day, J.J., Faircloth, B.C., Friedman, M. & Near, T.J. 2024 Dispersal sweepstakes: Biotic interchange propelled air-breathing fishes across the globe. Journal of Biogeography 51, 797–813.

[51] Greenwood, P.H., Rosen, D.E., Weitzman, S.H. & Myers, G.S. 1966 Phyletic studies of the teleostean fishes, with a provisional classification of living forms. Bulletin of the American Museum of Natural History 131, 339–456.

[52] Schaeffer, B. & Mangus, M. 1976 An Early Triassic fish assemblage from British Columbia. Bulletin of the American Museum of Natural History 156, 515–564.

